# Differential variation of NSCs in root branch orders of *Fraxinus mandshurica* Rupr. seedlings across different drought intensities and soil substrates

**DOI:** 10.1101/2021.04.08.439108

**Authors:** Li Ji, Yue Liu, Jun Wang, Zhimin Lu, Yuchun Yang, Lijie Zhang

## Abstract

Non-structural carbohydrates (NSCs) facilitate plants adapt to drought stress, could characterize trees growth and survival ability and buffer against external disturbances. Previous studies have focused on the distribution and dynamics of NSCs among different plant organs under drought conditions. However, discussion about the NSC levels of fine roots in different root branch order were little, especially the relationship between fine root trait variation and NSCs content. The aim of the study is to shed light into the synergistic variation of fine root traits and NSC content in different root branch order under different drought and soil substrate conditions. 2-year-old *Fraxinus mandshurica* Rupr. potted seedlings were planted in three different soil substrates (humus, loam and sandy-loam soil) and conducted to four drought intensities (CK, mild drought, moderate drought and severe drought) for two months. With the increase of drought intensity, the biomass of fine roots decreased significantly. Under the same drought intensity, seedlings in sandy-loam soil have higher root biomass, and the coefficient of variation of fifth-order roots (37.4%, 44.5% and 53.0% in humus, loam and sandy loam, respectively) is higher than that of lower-order roots. With the increase of drought intensity, the specific root length (SRL) and average diameter (AD) of all five orders increased and decreased, respectively. The fine roots in humus soil had higher soluble sugar content and lower starch content. Also, the soluble sugar and starch content of fine roots showed decreasing and increasing tendency respectively. Soluble sugar and starch explain the highest degree of total variation of fine root traits, that is 32.0% and 32.1% respectively. With ascending root order, the explanation of the variation of root traits by starch decreased (only 6.8% for fifth-order roots). The response of different root branch order fine root morphological traits of *F. mandshurica* seedlings to resource fluctuations ensures that plants maintain and constructure the root development by an economical way to obtain more resources.

## Introduction

In recent decades, forest decline and death caused by high temperatures and extreme droughts, have occurred on a large scale worldwide (Allen et al., 2010; Choat et al., 2012; Zhang et al., 2015; Martínez□Vilalta et al., 2016), and global climate change is predicted tree death is becoming more and more serious, which inevitably affect the carbon metabolism and balance in the plant and change its physiological metabolic function (Choat et al., 2012; Adams et al., 2013). Non-structural carbohydrates (NSCs), as important substances involved in the life process of plants, are mainly composed of soluble sugars and starches, they largely reflect the carbon supply status of plants and affect the growth and development of plants (Richardson et al., 2013). Moreover, the size of their content can characterize the buffering capacity to cope with the pressure of environmental stress (Xie et al., 2018). When plants undergone drought stress, the stored NSCs can be used as a buffer to temporarily supply plants for their growth and metabolism (Dietze et al., 2014). In recent years, the different responses of NSCs among different plant tissues or organs under drought stress were discussed deeply (Martínez□Vilalta et al., 2016; Furze et al., 2019; Deng et al., 2020; He et al., 2020; Zhang et al., 2020). Many studies have found that roots have the highest concentration of NSC except for trunk, which had an important impact on NSC storage and distribution in trees (Dietze et al., 2014; Mei et al., 2015; Ji et al., 2020). However, there have been few empirical investigations into the variation of fine roots NSC (especially the functional root branch order) in response to drought. Therefore, understanding the variation of the composition and level of fine root NSC under drought conditions has great significance for better recognition of the carbon balance and dynamics of plant survival and growth (Hartmann and Trumbore, 2016).

Root plays a crucial role in plant growth and productivity, especially in resource-constrained environments. The morphological and physiological plasticity of root reflect an important mechanism for plants to obtain limited soil resources (Ristova and Busch, 2014; Rogers and Benfey, 2015). Fine roots (≤2 mm) are the main organ for water and nutrient absorption, and the most active and sensitive part of the root system (McCormack et al., 2015; Ma et al., 2018). Many scholars pointed out that changes in root starch under drought conditions are related to plant survival, and root NSCs reserves play an important role in repairing embolism and preventing self-death (Rodríguez□Calcerrada et al., 2017; Kannenberg et al., 2018). Oswald and Aubrey (2020) observed that the root starch concentration of *Pinus palustris* in xeric environments increased delayed in summer compared with mesic habitats. Starch stored in the root system can promote root growth and maintain root osmotic potential, ensuring plants can absorb more water (Ge et al., 2012; Camarero et al., 2016). However, previous studies have mostly focused on the overall NSCs level of the root system (including coarse roots and fine roots) (Hoch and Körner, 2003; Landhäusser and Lieffers, 2012; Hartmann, 2013; Yang et al., 2021), nevertheless, the response of fine root NSCs with functional root branch orders to drought is rarely involved (Aubrey and Teskey, 2018; Nikolova et al., 2020). Pregitzer et al. (2002) found that the lower-order roots were produced later, the younger the age, the larger the contribution, the lower-order roots have higher nitrogen content and the respiration metabolism ability, and they are more sensitive and fragile than the higher-order roots. Guo et al. (2008b) conducted a study on the root for twenty-three tree species in the temperate of China, they pointed out that the first three roots of trees are generally non-lignified, with primary structures and complete cortical tissues, and their mycorrhizal colonization rate is relatively higher, and they mainly perform the function of nutrient and water absorption. most of the fourth- and fifth-orders roots have been lignified, and the cortex tissue has disappeared, with a continuous cork layer and secondary xylem, which mainly play a role of transport and storage. In addition, previous studies mostly compared the effects of single drought or severe drought on the root system, but far too little attention had been paid to the gradient studies (Hartmann et al., 2013; Kannenberg et al., 2018; Blackman et al., 2019; Zhang et al., 2020). McDowell et al. (2008) speculated that trees only depleted NSC under mild or moderate drought, while severe drought will cause the xylem to form embolism while NSC is not depleted. Recently, a meta-analysis based on fifty-two tree species around the world indicated that variations in plant NSCs were related to drought intensity, and the net loss of carbohydrates from roots is the most obvious (He et al., 2020). Therefore, this indicates a necessary to shed light into the response of fine root NSCs exist among different root orders across drought intensity.

Root traits play a vital role in the acquisition and transportation of water and nutrients. Therefore, they can strongly affect plant growth, survival and response to climate change (Bardgett et al., 2014; Kong et al., 2014; Iversen et al., 2017). Compared with leaf traits, the response of fine root traits to environmental changes can reflect adaptation strategies for resource utilization and plant performance under climate change (Bardgett et al., 2014; Warren et al., 2015). Moreover, root traits have greater variability and uncertainty (Comas and Eissenstat, 2004). Although numerous studies have reported the response of root traits to drought or water deficit (Comas et al., 2013; Fort et al., 2017; Zhou et al., 2018; Zhou et al., 2019;Lozano et al., 2020;Nikolova et al., 2020), few studies have focused on the synergistic changes and relationships between root NSCs levels and root traits under stress condition (Ji et al., 2020; Yang et al., 2021). Olmo et al. (2014) studied the drought resistance response of ten woody tree species seedlings and found that SRL increased significantly under drought conditions. It is a strategy that when water is limited, plants can build longer root with less carbon. Indeed, increasing carbon input per unit produces a larger surface area, length of fine roots and more fine roots, it could facilitate to optimize the cost-benefit ratio of fine roots (Eissenstat et al., 2000; Ostonen et al., 2007b). Under drought conditions, thicker roots with transport and storage functions tend to preserve NSC (Konôpka et al., 2007; Yang et al., 2021), while thinner roots with absorption function are severely affected by drought (Olmo et al., 2014). Yang et al. (2021) found that significant correlation between the roots NSC concentration, root architecture and SRL occurred in *Phyllostachys edulis* seedling under drought conditions, indicating that the sensitivity of NSC concentration to drought supported the plasticity of root architecture to a certain extent, and plants could build low-cost roots through more carbon investment. In addition, when plants adapt to drought stress, they balance the lack of tissue radial growth by increasing the concentration of NSC in the growing parts (Dietze et al., 2014; Kannenberg et al., 2018). Therefore, exploring the coupled relationship between the variation of fine root NSCs and root traits under drought conditions will help us to further understand the response strategies of plants to water deficit.

*Fraxinus mandshurica* Rupr. is one of the main timber species in northeastern China. It has been proven that its root system (shorter primary root and developed lateral roots) had a branch sequence of primary (non-woody) structure and developed into a branch through secondary development. The branched order of woody roots is sensitive to nutrients and water (Wei et al., 2009; Xia et al., 2010). In Jilin Province China, *F. mandshurica* plantations are distributed in north-south latitudes, and the soil types they inhabit are roughly divided into three categories, namely, humus, loam and sandy-loam soil. Weemstra et al. (2017) found that the specific root length (SRL) and root tissue density (RTD) of the fine roots of European beech and Norway spruce were not significantly different in clay and sandy soil, but the dry mass of fine roots in sandy (both species) was ten times than that in clay. Paudel et al. (2016) had studied these interactions in the osmometer of the orchard and the clay sandy loam, and found that soil type significantly affects the morphological traits of all root branch order of *Citrus paradisi* Macf. The interaction of root system with soil quality and water will cause changes in root growth, structure and function (Paudel et al., 2016). To our knowledge, few studies have focused on the response of fine root NSC to drought intensity gradient in different soil substrates, and even less is known regarding to the coupled relationship between fine root NSC and traits. In this study, two-year-old *F. mandshurica* potted seedlings with different drought intensities and soil substrates were set up, and compare the variation and the coupled relationship of the fine root traits and NSCs content of different root branch order of *F. mandshurica* seedlings under different water and soil conditions. We hypothesize that 1) with the increase of drought intensity, the specific root length of fine roots increases and the root diameter decreases; the specific root length in humus soil is the lowest and the root diameter is thicker; 2) with the increase of drought intensity, the variation of root carbon, nitrogen and phosphorus is slight, but the variation of NSC will be obvious, and the root soluble sugar content will decrease, and the starch content will increase; 3) root morphological traits are closely related to the NSCs content.

## Materials and Methods

### Experimental site and sapling preparation

A controlled pot experiment was conducted at the Xinli Town, Jingyue Development District, Changchun, which is located in Jilin Province, China (43°33′ N-44°41′ N, 125°19′ E-125°24′ E). The location belongs to temperate continental monsoon climate with a frost-free period of 140 days; and with mean annual rainfall of 600-800 mm, which mainly falls from July to September and a mean annual temperature of 4.6□.

The two-year-old *Fraxinus mandshurica* Rupr. seedlings of the Hongwei nursery of Lushuihe Forestry Bureau, in Jilin Province were used as experimental materials. They were transplanted into plastic pots (24 cm × 20 cm) in the April, 2017, and were placed in the flat, open canopy before they started to bud. The cultured soil in pots was filled with equal volumes of humus, loam and sandy-loam respectively, and the humus was collected in coniferous and broad-leaved forest in Lushuihe Forestry Bureau. The main soil type is an Eum-Orthic Anthrosol according to the Food and Agricultural Organization soil classification system. The soil surface was well ventilated throughout the experiment. At the experimental site, the pots were placed in rows with 50 cm apart from the neighbors under full sunlight. They were kept well-watered prior to the application of drought treatments, and the gravimetric soil water content was initially maintained at field capacity. Fertilizer was not added during the experiment period.

### Experimental design and sampling

Before the beginning of the drought experiment, three soil substrates samples were collected in July 2017 to determine soil total nitrogen, total phosphorus, available phosphorus and soil physical structure, water content and field water holding capacity. The one hundred-twenty pots of cultured *F. mandshurica* seedlings were randomly selected from three different substrates to conduct a water control experiments. The basal diameter and height of each seedlings were measured using a vernier caliper with an accuracy of 0.01 mm and a tape measure with an accuracy of 0.1 cm prior to the application of drought treatments in July 2017, respectively.

The soil substrates and drought were conducted for two-factor complete orthogonal design. Three soil substrates were set with four drought stress gradients, the control (CK): approximately 80∼85% of the maximum field water-holding capacity; mild drought (T1): 60%∼65%; moderate drought (T2): 40∼55%; and severe drought (T3): 20∼25%. The total of 12 treatments, each treatment set 3 blocks, selected 20 pots of seedlings in each block (selecting uniform seedlings). The detailed description of the drought control refers to our previous study (Ji et al., 2020).

In each experimental block, ten seedlings were randomly selected and the roots of the sampling were destructively sampled after two months of continuous drought stress. Root were sorted carefully out of the soil, and root samples were washed free of soil particles by deionized water until the branching structure of the roots can be identified, and then put it into the labeled pocket and store it in a freezer (2-3°C). Then divided the root sample into two parts: (1) root morphology analysis sample; (2) chemical characteristic analysis sample. All samples were shipped back to the laboratory on the same day and stored in a freezer at -20 °C. In this study, we only measured live root samples, and the dead roots were picked and discarded.

### Soil physicochemical property

The soil physical and chemical properties of the three soil substrates were determined before drought stress (Table 1). The soil total nitrogen was determined by Kjeldahl titration. The soil total phosphorus was determined by the sulfuric acid-perchloric acid-molybdenum anti-colorimetric method (Yang et al., 2018). Soil available phosphorus was extracted by double acid extraction, soil water content and bulk density were determined by ring knife method (Yang et al., 2018), soil total porosity, aeration porosity, water absorption multiple, water seepage rate and evaporation rate reference (Wei et al., 2015) method determination.

**Table 1.** Physicochemical properties of three soil substrates.

| Soil property | Humus | Loam | Sandy-Loam |
| --- | --- | --- | --- |
| Bulk density ( g·cm<sup>-3</sup> ) | 1.18±0.02b | 1.35±0.02a | 1.32±0.02a |
| Total porosity ( % ) | 53.19±1.12b | 58.90±0.91a | 36.72±0.95c |
| Aeration porosity ( % ) | 25.61±0.48a | 23.52±0.53a | 13.01±0.99b |
| Water absorption capacity | 0.23±0.01b | 0.26±0.01a | 0.18±0.01c |
| Penetrate rate(g·min<sup>-1</sup>) | 4.49±0.04a | 4.01±0.12b | 3.04±0.18c |
| Evaporation rate(g·h<sup>-1</sup>) | 0.59±0.01a | 0.50±0.01b | 0.37±0.01c |
| Total nitrogen (mg·g<sup>-1</sup>) | 7.09±0.78a | 3.11±0.05b | 1.20±0.03c |
| Total phosphorus (mg·g<sup>-1</sup>) | 0.71±0.04a | 0.34±0.09b | 0.32±0.01b |
| Available phosphorus (mg·kg<sup>-1</sup>) | 13.10±0.82a | 5.04±0.21b | 12.75±0.69a |

Note: The different letters in the same line indicate significant difference among the different treatments ( $P<0.05$ ).

### Fine root morphological and chemical traits

In the laboratory, root samples for morphological analysis were carefully dissected with forceps on the basis of branch order, following the procedure described in Pregitzer et al. (2002) and Wang et al. (2006), with the distal nonwoody roots regarded as frst-order roots (1st-order root), and the next root segment is the 2nd-order roots. Then, root samples were scanned with an Expression 10000XL 1.0 scanner in Northeast Forestry University (Epson Telford Ltd, Telford, UK). The mean diameter, total length and volume of root tips were determined with the root system analyzer software (WinRhizo 2004b, Regent Instruments, Inc., Québec, Canada). These root samples were oven-dried at 65°C to determine constant weight (nearest 0.0001 g) and calculated the specific root length (SRL), specific root surface area (SRA)and root tissue density (RTD).

For root chemical analyses, as mentioned above, the fine root sample after scanning was placed in a 65 °C and dried for 48 h to constant weight. The dried fine root sample was ground and homogenized by using a Ball mill instrument (RETSCH MM 400, Germany), and 1.00 g of the dry powder sample was weighed and pressed with a FYD-20 electric tableting machine to a boat which was thickness of 6 mm and a diameter of about 13 mm. For the pellet sample, the tableting conditions can be adjusted according to the actual conditions. The tableting conditions of the test are maintained at a pressure of 16 MPa for 3 min. The carbon, nitrogen and phosphorus element in the root samples after tableting was measured by a J200 Tandem laser spectroscopic element analyzer.

### Fine root NSCs concentration

NSC concentration was defined as the sum of soluble sugar (SS) and starch (ST) concentrations that were measured using the anthrone method (Yemm and Willis, 1954). Root sample (0.1000 g) was placed into a 10 ml centrifuge tube, and 2 ml of 80 % ethanol was then added. The mixture was incubated at 80□ in a shaking water bath for 30 min and then centrifuged at 4000 rpm for 5 min. A further two extractions from the pellets were carried out with 80% ethanol. The supernatant was retained, combined, and stored at -20□for soluble sugar determination.

Starch was extracted from the ethanol-insoluble pellet after ethanol was first removed by evaporation. The starch in the residue was then released by boiling in 2 ml distilled water for 15 min. After cooling to room temperature, 2 ml 9.2 M HClO_4_ was added, and the mixture was shaken for 15 min. Four milliliters of distilled water were then added, and the mixture was centrifuged at 4000 rpm for 5 min. A further extraction was carried out with 2 ml 4.6 M HClO_4_. The supernatant was also retained, combined, and stored at -20□for starch determination.

Soluble sugar and starch determination were performed based on the absorbance at 625 nm using the same anthrone reagent in a spectrophotometer (Yemm and Willis, 1954). Sugar concentration was calculated from the regression equations based on glucose standard solutions and starch concentration by multiplying glucose concentration with a conversion factor of 0.9 (Osaki et al. 1991).

### Data analysis

Normality and variance homogeneity requirements were met, and no data transformation was necessary. The Three-way ANOVA analysis of fine root traits and NSC content was performed by SPSS19.0 (IBM Co., Armonk, NY, USA) and examine the differences in fine root traits of seedlings between different treatments (LSD, α=0.05); Principal component analysis (PCA) and redundancy analysis (RDA) were performed on the fine root traits of seedlings under different treatments using Canoco software (Version 4.56, Biometris Plant Research International Wageningen, The Netherlands). The Monte Carlo test was performed on the parameters in the RDA analysis using the R software (vegan package) (R Core Team, 2018). A multiple linear regression analysis was done using the Sigmaplot 12.5 software (Systat Software Inc., San Jose, California, USA) to analyze the influence of root traits on the fine root NSCs (soluble sugar and starch) content in all treatments. All data are mean ± standard error (Mean ± SE). All bar figures were drawn using Origin Pro 8.5 (OriginLab, Northampton, MA, USA).

## Results

### Fine root biomass and morphological traits among root branch order

Soil substrates, drought intensity and root order had significant differences in the fine root biomass of *F. mandshurica* seedlings (Table S1). With the increase of drought intensity, the fine roots biomass of seedling in the three substrates showed a decrease progressively (Figure 1A∼1E). The biomass of fifth-order roots had the highest variation, which was 37.4%, 44.5% and 53.0%in humus, loam and sandy loam, respectively (Figure 1E). Under the same drought treatment, the fine root biomass of seedlings for all branch order was the highest in sandy-loam soil and the lowest in humus soil. With the increase of drought intensity, the coefficient of variation in the fifth-order root biomass among different soil substrates was the highest, which were 61.8% (in CK), 36.6% (in T1), 57.0% (in T2) and 40.4% (in T3) (Figure 1E).

**Figure 1.**
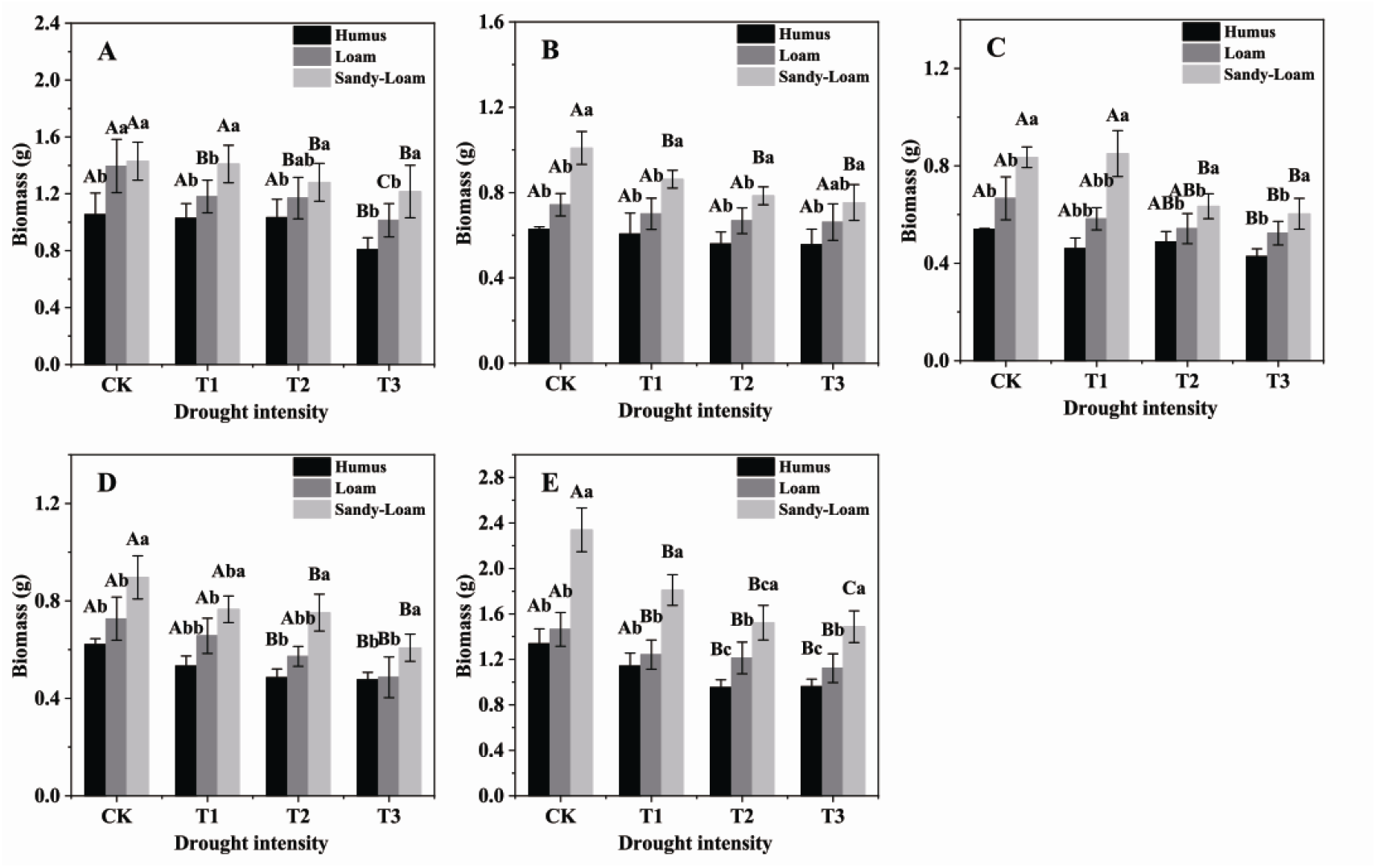
The fine root biomass of *F. mandshurica* seedlings in different drought intensities and soil substrates. The different lowercase letters denote significant differences among soil substrates (*P* <0.05). The different uppercase letters denote significant differences among drought intensities (*P* <0.05). **(A)**, 1st order root; **(B)**, 2nd order root; **(C)**, 3rd order root; **(D)**, 4th order root; **(E)**, 5th order root. CK, control; T1, mild drought; T2, moderate drought; T3, severe drought.

Soil substrate, drought intensity and root order had significant effects on the specific root length (SRL), specific root surface area (SRA) and root tissue density (RTD) of *F. mandshurica* seedlings (Table S1). Under the same drought intensity, the SRL and SRA of all branch order in the humus soil were the lowest, and showed a significant decrease with ascending root order (Figure 2A∼2C, Figure 2D∼2F). The average diameter (AD) and RTD of the fine roots of all branch order were the highest and lowest in the humus and sandy-loam soil, respectively, and showed a significant increase trend with ascending root order (Figure 2G∼4I, Figure 2J∼5L). With the increase of drought intensity, the SRL and SRA of seedlings for all soil substrates increased significantly. Compared with CK, the SRL and SRA under T3 treatment increased significantly by 39.0% (variation range 19.1%∼88.6%) and 22.3% (variation range 9.9%∼37.4%). Compared with CK, the RTD of seedlings in all the three soil substrates was the lowest under T3 treatment, especially in the 1st root order, which decreased by 13.7% (humus soil), 10.7% (loam soil) and 28.6% (sandy-loam) respectively (Figure 2J). The AD of all root branch order was less affected by drought intensity (Figure 2G∼2I, Table S1).

**Figure 2.**
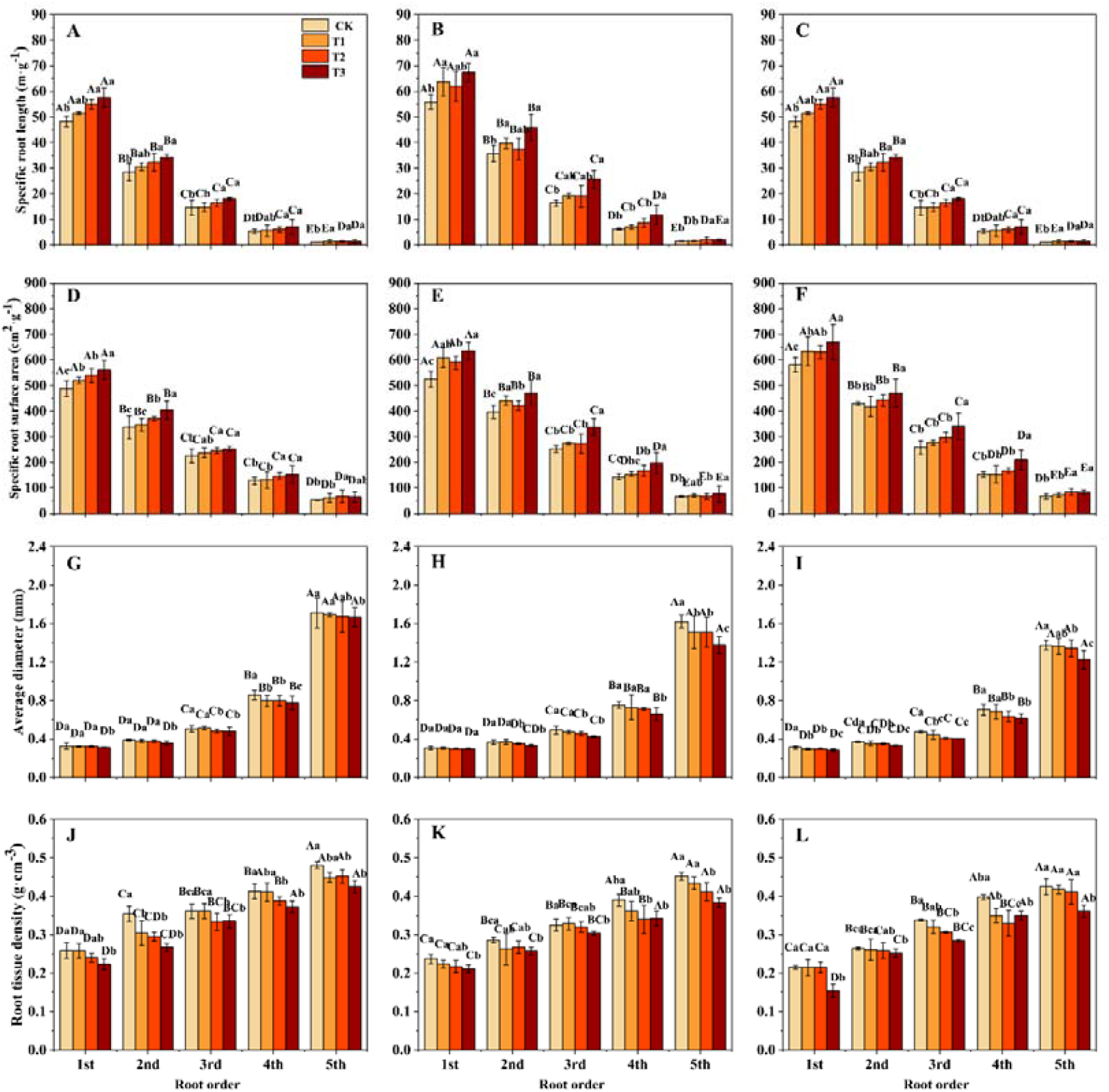
The specific root length **(A, B, C)**, specific root surface area **(D, E, F)**, average diameter **(G, H, I)**, root tissue density **(J, K, L)** of *F. mandshurica* seedlings in different drought intensities and soil substrates. The different lowercase letters denote significant differences among drought intensities (*P* <0.05). The different uppercase letters denote significant differences among root branch order (*P* <0.05). **(A, D, G, J)**, humus soil; **(B, E, H, K)**, loam soil; **(C, F, I, L)**, sandy-loam soil; CK, control; T1, mild drought; T2, moderate drought; T3, severe drought.

### Fine root chemical traits among root branch order

The fine root carbon, nitrogen and phosphorus contents of *F. mandshurica* seedlings were significantly different among root branch orders. With ascending the root orders, the fine root carbon and nitrogen content showed a decreasing trend, while the fine root phosphorus content showed an increasing trend (Table S2, Figure 3A∼3C and 4A∼4C). The soil substrate had a significant effect on the fine root carbon and nitrogen content of *F. mandshurica* seedlings (Table S2). The carbon and nitrogen content of all branch orders in sandy-loam soil were significantly higher than those of humus soil. The carbon and nitrogen content of 1st-order root in sandy-loam soil was highest, which was 41.21 mg·g^−1^ and 2.14 mg·g^−1^, respectively (Figure 3A, 3B). Drought had a significant effect on the fine root carbon and phosphorus content of *F. mandshurica* seedlings, and the phosphorus content of first forth root order under T3 treatment was lower than that of CK (Figure 4C).

**Figure 3.**
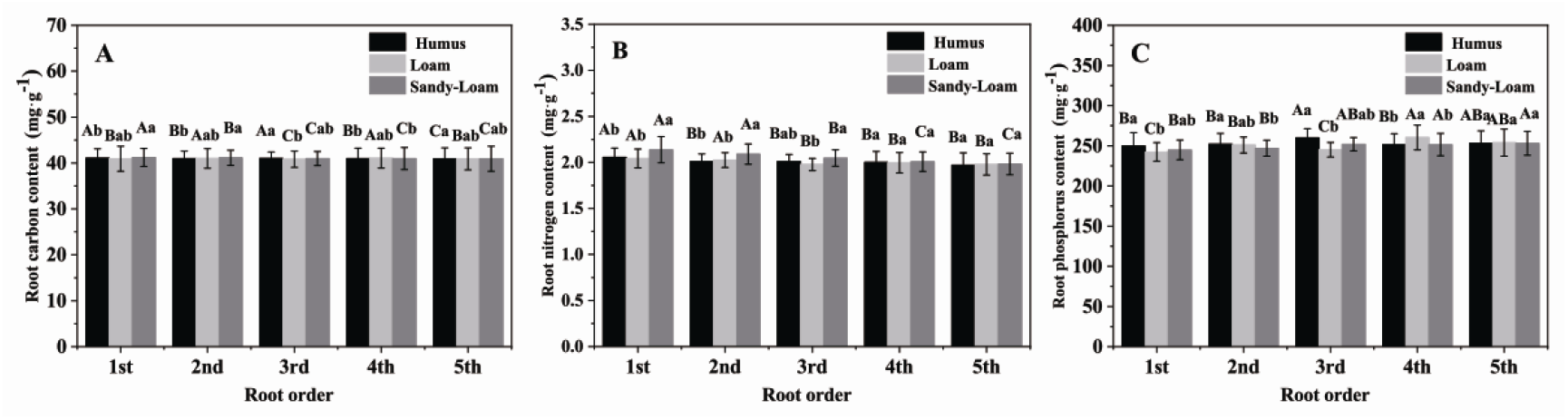
The root carbon **(A)**, nitrogen **(B)** and phosphorus **(C)** of *F. mandshurica* seedlings in different soil substrates. The different lowercase letters denote significant differences among soil substrates (*P* <0.05). The different uppercase letters denote significant differences among root branch order (*P* <0.05).

**Figure 4.**
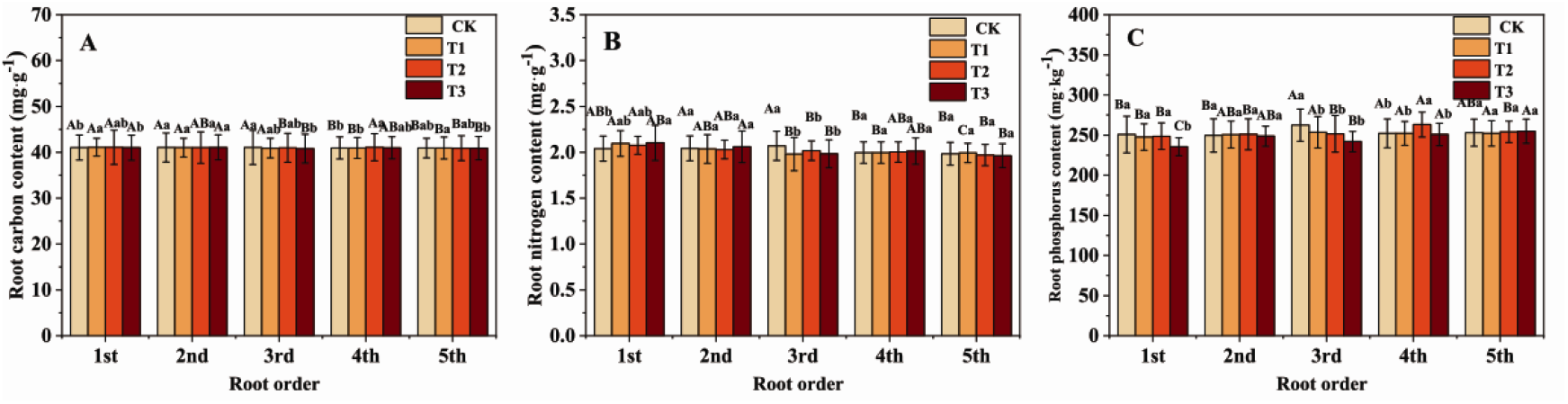
The root carbon **(A)**, nitrogen **(B)** and phosphorus **(C)** of *F. mandshurica* seedlings in different drought intensity. The different lowercase letters denote significant differences among drought intensities (*P* <0.05). The different uppercase letters denote significant differences among root branch order (*P* <0.05).

### Fine root NSCs content among root branch order

Soil substrate, drought intensity and root order had significant effects on the soluble sugar (SS), starch (ST) and total NSC content of the fine roots of *F. mandshurica* seedlings (Table S3). Under the same drought intensity, the SS content of seedlings in humus soil was the highest for all branch orders. With ascending root order, the SS content of fine roots of seedlings in humus soil was 107.9%, 162.7%, 125.7%, 269.2% and 118.5% higher than those of sandy-loam soil, respectively (after the average the intensity of the four droughts) (Figure 5A∼5E). For all soil substrates, the SS content of the fine roots of seedlings decreased with the increased drought intensity. With ascending root orders, the SS content of fine roots under T3 treatment was 51.3%, 58.1%, 62.1%, 68.7%, and 36.5% lower than those of CK, respectively (after the average of the three soil substrates) (Figure 5A∼5E). Under the same drought intensity, seedlings in sandy-loam soil had the highest starch and total NSC content for all root orders. The starch and total NSC content of lower-order roots in different soil substrates varied greatly. The starch and total NSC content of the 1st-order and 2nd-order roots in sandy-loam soil were 276.1%, 231.1%, 195.8% and 186.7% higher than those of humus soil respectively (after averaging the intensity of the four droughts) (Figure 5A∼5E). For all soil substrates, the fine root starch content generally increased with the increase of drought intensity. The starch and total NSC content of the 1st-order roots of seedlings under T3 treatment were 58.6% and 43.4% higher than those of CK, respectively (Figure 5A).

**Figure 5.**
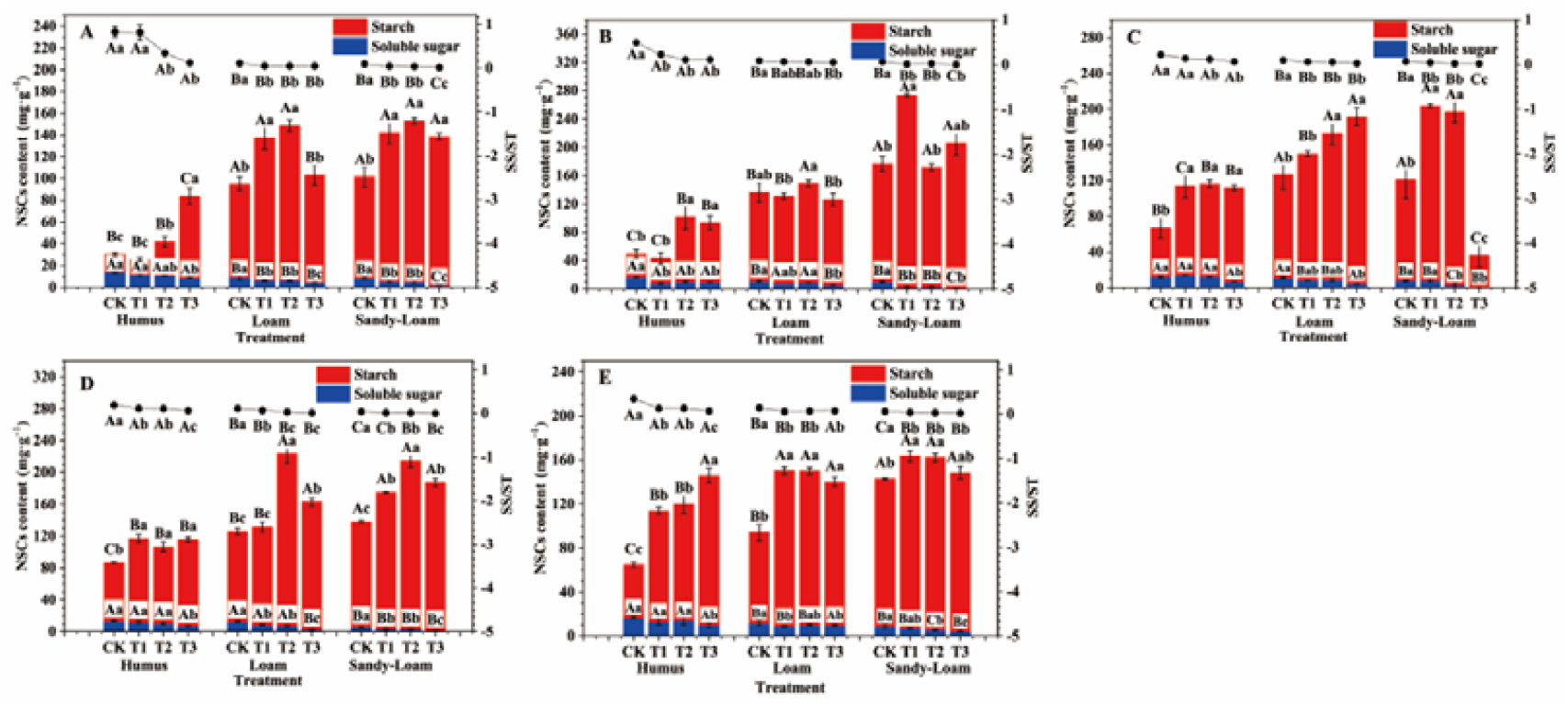
The fine root NSCs content of *F. mandshurica* seedlings in different drought intensities and soil substrates. The histogram represents the soluble sugar and starch content, and the line chart represents the soluble sugar-starch ratio (SS/ST). The different lowercase letters denote significant differences among soil substrates (*P* <0.05). The different uppercase letters denote significant differences among drought intensities (*P* <0.05). **(A)**, 1st order root; **(B)**, 2nd order root; **(C)**, 3rd order root; **(D)**, 4th order root; **(E)**, 5th order root. CK, control; T1, mild drought; T2, moderate drought; T3, severe drought.

### Relationship among the fine root biomass, traits and NSCs content

The root morphological and chemical traits of first five order root of *F. mandshurica* seedlings under different drought intensities and soil conditions were analyzed by redundancy analysis (RDA), the results showed that the first two axes of the RDA explained approximately 65% of the total variations between all treatments (Figure 6A∼6E). The first and the second ordination axis indicated the variations of fine roots morphological traits, and chemical traits and biomass, respectively. Soil substrates and drought intensity in plots had a good degree of separation. The fine roots morphological and chemical traits in all conditions were conducted partial Monte Carlo test. For the 1st-order roots, SS and ST explained the highest degree of total variation of fine root traits, which were 32.0% and 32.1%, respectively (Table S4). With ascending root orders, the explanation of the variation of root traits by starch decreased, only 6.8% (for 5th-order root) (Table S4).

**Figure 6.**
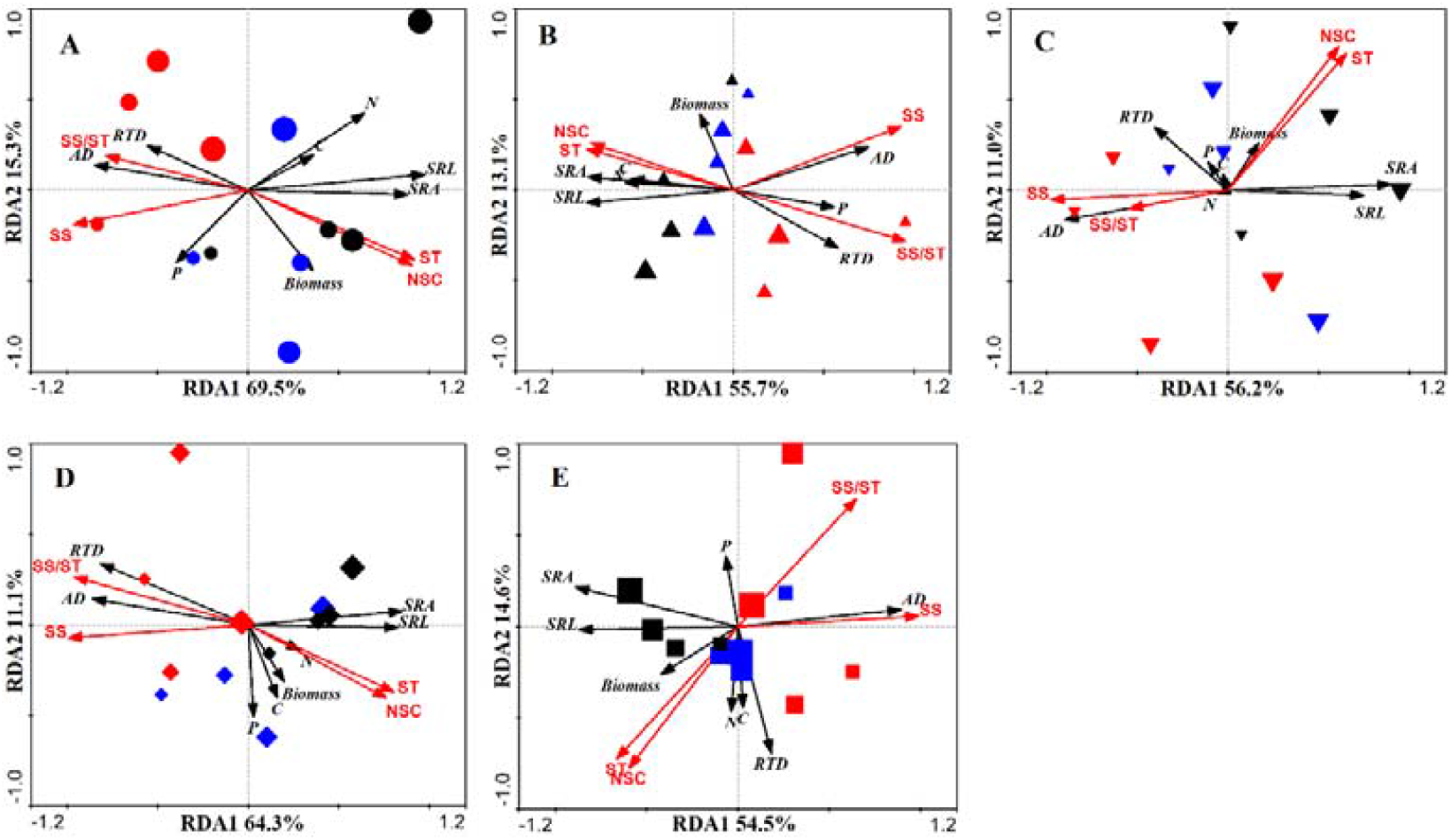
Redundancy analysis of fine root morphological and chemical traits in different drought intensity and soil substrates. **(A)**, 1st-order root; **(B)**, 2nd-order root; **(C)**, 3rd-order root; **(D)**, 4th-order root; **(E)**, 5th-order root. Red symbols, humus; Blue symbols, loam; Black symbols, sandy-loam. SRL, specific root length; SRA, specific root surface area; AD, average diameter; RTD, root tissue density; SS, soluble sugar; ST, starch; C, root carbon; N, root nitrogen; P, root phosphorus.

The Pearson correlation analysis showed that the first five order root morphological and chemical traits were significantly correlated with SS, SRL, SRA and root nitrogen content are significantly negatively correlated with SS, while the AD, RTD and root phosphorus content were significantly positively correlated with SS (Figure 7A∼7D).

**Figure 7.**
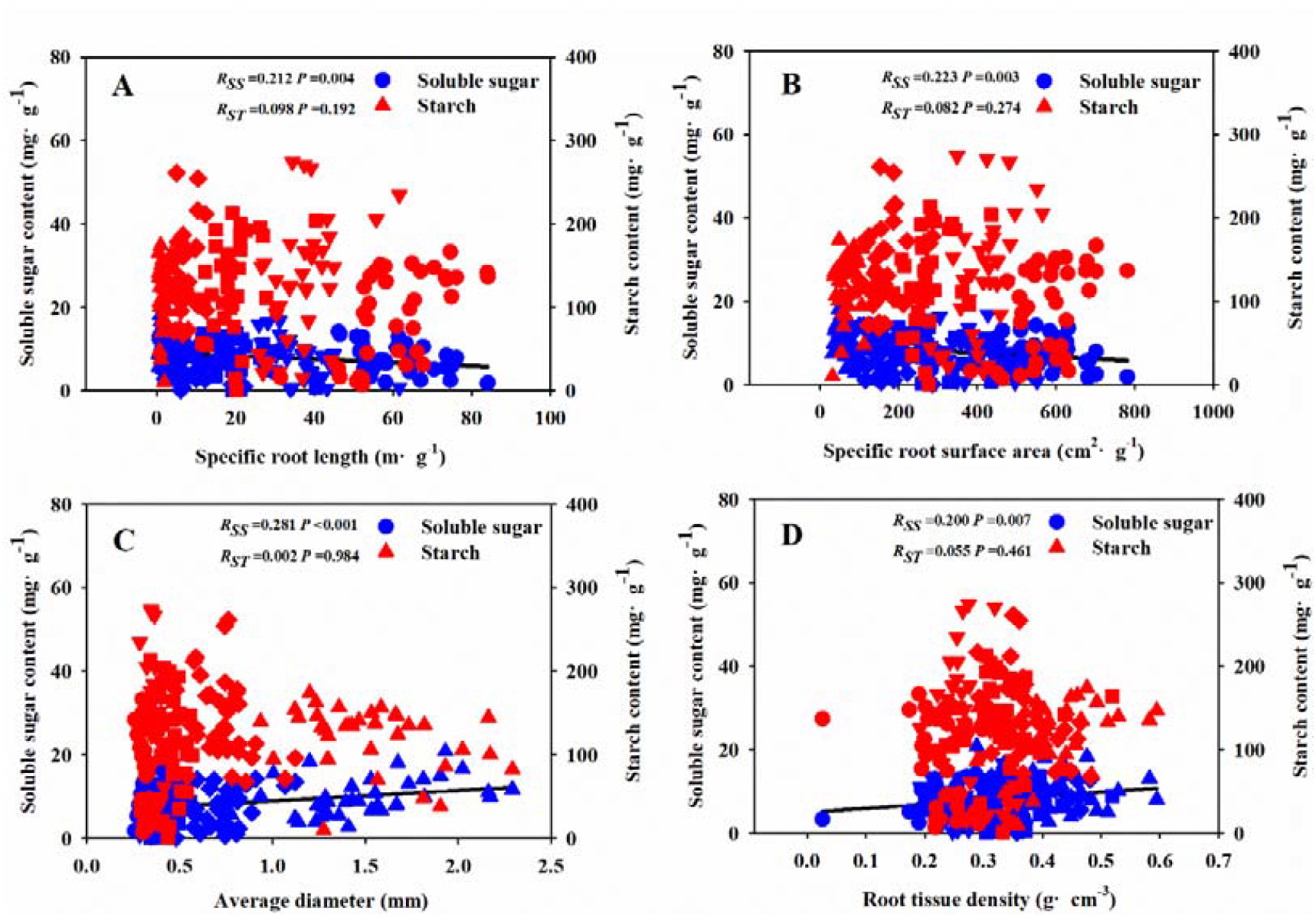
Relationships between soluble sugar, starch concentration and fine root morphological traits (SRL **(A)**, SRA **(B)**, AD **(C)** and RTD **(D)**) during the whole experiment period. R: correlation coefficient. SS, soluble sugar; ST, starch; circle represents 1st-order root; down-triangle represents 2nd-order root; square represents 3rd-order root; diamond represents 4th-order root; up-triangle represents 5th-order root;

## Discussion

Our results highlighted several key findings related to NSC and fine root traits of *F. mandshurica* seedlings under different soil substrates and drought intensities. Firstly, with the increase of drought intensity, the fine roots biomass decreased significantly. Under the same drought intensity, there had a higher biomass in relatively poor sandy-loam soils, and the coefficient of variation for fifth-order roots was higher than those of lower-order roots. Secondly, compared with chemical traits, fine root morphological traits were more sensitive to soil substrates and drought intensities. With the increase of drought intensity, the SRL and AD of all root orders increased and decreased respectively. Finally, the fine roots in the humus soil had higher soluble sugar content and lower starch content. With the increase of drought intensity, the soluble sugar and starch content of the fine roots showed a decreasing and increasing trend, and the NSC content of the fine roots significantly related to root morphological traits. According to the rapid response of root morphology to changes in the habitat environment, plants will maintain and constructure the development of underground organs in an economical way to obtain more resources and nutrients.

### Response of root morphology and biomass to drought and soil substrate

In our study, the fine root biomass of seedlings in sandy-loam soil (soil nutrient relatively poor) was significantly higher than that in humus soil (soil nutrient relatively rich), which was in line with those of previous studies (Hertel et al., 2013; Poorter and Ryser, 2015). Weemstra et al. (2017) pointed out that the fine-root mass and the growth rate of fine roots of both species of *Fagus sylvatica* and *Picea abies* in sandy soil were three times and ten times higher than those of clay soil, despite the family of tree species planted in different soils are not consistent. Hendricks et al. (1993) proposed two hypotheses: 1) As nitrogen availability increases, the carbon allocated to fine roots decreases, and the lifespan or turnover rate of fine roots remains unchanged; 2) as nitrogen availability increases, the distribution to the carbon in fine roots remains unchanged, the turnover rate increases, and the fine roots biomass decreases. Both hypotheses believed that the increase in soil nutrient availability will lead to a decrease in the fine root biomass. Liu et al. (2020) found that the higher-order root biomass of *Pinus tabuliformis* seedlings under the 20% field water holding capacity treatment was 33.3% lower than that under the 80% field water holding capacity treatment. We found that the biomass of higher-order roots (e.g., fifth-order) has the highest coefficient of variation under different soil substrates and drought treatments. This result may be explained by the fact that the different responses of fine roots of different root branch order to resource changes (Withington et al., 2006; Guo et al., 2008a). Indeed, with ascending the root order, the diameter of fine roots increased significantly (Pregitzer et al., 2002), and higher-order roots have a large number of wooden vessels that could transport more nutrients to the aboveground parts (Chapin III et al., 1990; Hishi and Takeda, 2005), and lower-order roots were mainly responsible for the absorption of water and nutrients, and were active in their growth and elongation (Chapin III et al., 1990; Noguchi et al., 2013).

The morphological traits of fine roots, such as AD, SRL, and RTD, are important functional parameters that characterize or affect the water absorption efficiency and ability of roots. In general, fine roots with smaller diameter and larger root length have higher water absorption efficiency (Freschet and Roumet, 2017; Dhiman et al., 2018; Ma et al., 2018). Root morphology characteristics responded to drought here in accordance with our first hypothesis as well, that was the root system of seedlings in humus soil had lower SRL, lower SRA, higher AD and RTD, these results were likely to be related to the higher nutrient content in humus soil, which matched those observed in earlier studies on the fine root traits of *Pinus tabuliformis, Larix gmelinii* and *F. mandshurica* after nutrient availability increased (Liu et al., 2009; Wang et al., 2013). Gruber et al. (2013) indicated that soil nitrogen deficiency usually promoted the elongation of main roots and some lateral roots. Plants can increase the absorption capacity of the root system in two different ways to adapt to water or nutrient shortages, 1) increase root yield and maintain a larger absorption surface area (acquisition strategy), or 2) increase the efficiency per unit mass absorption by changing the morphology and physiological condition of the root system (conservative strategy) (Lõhmus et al., 2006; Ostonen et al., 2007a). In our study, with the increase of drought intensity, the SRL and SRA of first five order roots increased significantly, and the RTD decreased significantly, which was a strategy to water deficit conditions of *F. mandshurica* seedlings. In addition, our results showed that the difference in various morphological trats among root orders had reached a significant level, indicating that the fine roots of *F. mandshurica* had a high degree of morphological heterogeneity, which was consistent with the previous study of the plasticity of the morphological traits of the root system from the perspective of orders (Pregitzer et al., 1997; Pregitzer et al., 2002; Wang et al., 2006). Cortina et al. (2008) found that *Pistacia lentiscus* seedlings could rapidly expand the length and surface area of fine roots when exposed to drought stress, and shaped the fine roots to avoid the damage of arid environment. In general, the larger the SRL or SRA and the smaller the AD, the higher the water absorption efficiency of fine roots (Freschet and Roumet, 2017; Dhiman et al., 2018). Our results emphasized the adaptation strategies of the fine roots of *F. mandshurica* seedlings in three soil substrates to different drought intensities.

### Effects of drought and soil substrate on root NSC

Soluble sugar is an important osmotic adjustment substance for plants to tolerate arid environment. It could reflect the drought status of plants. The variation of its concentration changes can adjust the osmotic pressure of cells in plants to maintain normal physiological activities to adapt to drought stress (Quentin et al., 2015). In this study, compared with sandy-loam soil, the soluble sugar content of fine roots (especially lower-order roots) in humus soil significantly increased, the starch content significantly decreased, the NSC significantly decreased, and the soluble sugar-starch ratio increased significantly. The development of fine roots accelerates the consumption of NSC, and the roots increase the ratio of soluble sugar-starch ratio to cope with stress conditions. The increase in the ratio is beneficial for plants to adjust osmotic potential to maintain the transport channel between leaves and roots, and improve water transport efficiency (Sala et al., 2012). Some studies had pointed out that lower-order roots might obtain greater carbon investment than higher-order roots, because the main function of lower-order roots is to absorb water and nutrients (absorptive root), while higher-order roots are responsible for transporting nutrients and supporting the entire root (Liu et al., 2020). Our results showed that after two months period of drought, the lower-order roots of *F. mandshurica* seedlings had more NSC content than the higher-order roots. This result implied that when carbon was limited, *F. mandshurica* preferentially allocate carbon to thin roots rather than thick roots. It supported the results of studies in hybrid poplar *Populus×canadensis* cv. *Eugeneii* (Kosola et al., 2001) and *Pinus tabuliformis* (Liu et al., 2020). Under carbon-limiting conditions, plants give priority to the distribution for fine roots. On the one hand, they could reduce carbon consumption (higher-order roots consume more carbon than lower-order roots). On the other hand, maintaining fine roots to absorb more water and nutrient (Finér et al., 2011). Liu et al. (2020) conducted the short-term drought stress on *Pinus tabuliformis* seedlings and found that the amount of ^13^C allocated to the first three roots 120 days after isotope labeling was significantly higher than that of the control in the moderate and mild drought, while the amount of ^13^C allocated to fifth-order roots was significantly higher in the control. Some studies had suggested that carbon starvation caused by drought stress may only exist in the belowground organ and had little to do with the aboveground part. Specially, thick roots are essential to alleviate the decline in the NSC content of the entire plant caused by drought (Hartmann, 2013; Kannenberg et al., 2018), drought will cause the loss of phloem function, making the aboveground and underground parts uncoupled. When plant is subject to drought stress, the NSC content of the aboveground organs will increase or remain stable for a short time, while the NSC content of the root will decrease significantly (Sevanto et al., 2014).

Galvez et al. (2011) found that drought significantly increased the concentration of soluble sugar and starch in the fine roots of *Populus tremuloides* seedlings. Our results supported the second hypothesis, that was with the increase of drought intensity, the variation of root NSCs was higher than that of root chemical traits, root SS content decreased, starch and total NSC content increased. A possible explanation for this might be that the path of NSC to the fine roots of *F. mandshurica* seedlings was blocked after two months drought stress, causing the carbohydrates produced by leaf photosynthesis to not be transported to the belowground organ. The fine roots could only rely on regulating their own NSC levels to deal with drought stress (Dietze et al., 2014). However, our inference needs to be further verified by combining the dynamic changes of fine root NSC content under different periods of drought treatment. Our results revealed the interaction of different soil substrates and drought intensities on the variations of first five root orders NSCs of *F. mandshurica* seedlings.

### Linking the fine root traits to the NSCs level

The root system is the organ responsible for absorbing water, and is the first responder to various stresses. Given that the root tends to grow in the moist soil, it can minimize the impact of water shortage (Brunner et al., 2015; Weemstra et al., 2016). Under drought stress conditions, more carbohydrates are allocated to lower-order roots to promote root structure and growth (Liu et al., 2020). Despite there is evidence that thick roots increase NSC accumulation under drought conditions (Yang et al., 2016), whether the variation of fine root traits is directly related to NSC accumulation remains to be explored. Our previous research pointed out that the root tip length and diameter of the seedlings (three broad-leaved tree species in the temperate zone) under drought conditions explained a higher variation of starch and soluble sugar (Ji et al., 2020). In this study, drought reduced the root biomass of *F. mandshurica* seedlings, changed the root traits of all root orders, and promoted the absorption of water and nutrients. This response may be related to the increased demand for osmotic active C compounds under drought conditions. NSC can provide fuel for root respiration and is an important substrate for root growth and physiological regulation (George et al., 2003; Xu et al., 2008). In this study, SRL and SRA of all samples were significantly negatively correlated with the soluble sugar content of fine roots, and AD and RTD were significantly positively correlated with starch content, which confirms our third hypothesis. When the diameter is smaller, the level of NSC required to build and maintain a thin root per unit length of the plant would be lower than that of the thicker root (Guo et al., 2004). Ma et al. (2018) pointed out that the root diameter of herbaceous plants were thinner than woody plants, so more carbon could be easily distributed to the roots under drought conditions. Maguire and Kobe (2015) pointed out that drought stress may directly increase root mortality by depleting starch and sugar reserves, and indirectly inhibited the transport of photosynthetic products to roots (Hasibeder et al., 2014). Therefore, our results suggested that changes in NSCs caused by changes in environmental conditions were related to the variation of fine root morphological traits.

## Conclusion

This project is the first comprehensive investigation of the interactive response of root branch order and fine root NSCs levels of *F. mandshurica* seedlings to drought intensity and soil substrate. Our results confirm that the variation of higher-order fine root biomass was higher than that of lower-order roots under different soil substrates and drought intensities. Secondly, the fine roots of the seedlings in the humus soil (first fifth root orders) have higher soluble sugar content and lower starch content. With the increase of drought intensity, the soluble sugar and starch contents of the fine roots showed decrease and increase trends respectively. The variation of fine root NSCs content was related to the variation of root morphological traits (SRL, SRA, AD, and RTD) induced by drought and soil substrate, rather than root chemical traits. This study reveals the adaptation strategies of *F. mandshurica* seedlings to drought under different soil substrate conditions, thereby enhancing the understanding of the construction and maintenance of the root system of *F. mandshurica*, and contributing to optimize soil water management in *F. mandshurica* plantations. In further research, it is necessary to combine ^13^C isotope labeling technology to more deeply reveal the mechanism of carbohydrate distribution among different root branch orders in a prolonged periods drought.

## Author Contributions

LJ and YY conceptualized the main question. LJ, JW and ZL conducted the field work. LJ and YL collected data and performed the data analyses. LJ wrote the manuscript. LJ, YY and LZ revised the manuscript. All authors read and approved the manuscript.

## Acknowledgments

We gratefully acknowledge the managers and workers of nursery for their logistic assistance with this project. We sincerely thank Wanying Cui, Sijia Liu, Jianhua Bi and Linlin Cao for helping with field work.

