## Supplemental files for "Differential variation of NSCs in root branch orders of *Fraxinus mandshurica* Rupr. seedlings across different drought intensities and soil substrates"

Supplementary Material

**Table S1** Results of three-way (drought intensity × soil substrate × root order) ANOVA of roots biomass and root traits. Values in bold type indicate significant effects

**Table S2** Carbon, nitrogen and phosphorus contents among root order of *F. mandshurica* seedlings and results of three-way (drought intensity × soil substrate × root order) ANOVA of root chemical traits.

**Table S3** Results of three-way (drought intensity × soil substrate × root order) ANOVA of roots NSCs content. Values in bold type indicate significant effects

**Table S4** Mantel analysis on the relationship between the root traits and NSC variables.

**Table S1** Results of three-way (drought intensity × soil substrate × root order) ANOVA of roots biomass and root traits. Values in bold type indicate significant effects

| Source of variation | df | *P* values | | | | |
| --- | --- | --- | --- | --- | --- | --- |
|  |  | Biomass | SRL | SRA | AD | RTD |
| Drought intensity (D) | 3 | **0.025** | **<0.001** | **<0.001** | 0.458 | **0.014** |
| Soil substrate (S) | 2 | **<0.001** | **<0.001** | **<0.001** | **0.002** | **0.016** |
| Root order (R) | 4 | **<0.001** | **<0.001** | **<0.001** | **<0.001** | **<0.001** |
| D×S | 6 | 0.956 | 0.803 | 0.618 | 0.998 | 0.159 |
| D×R | 12 | 0.991 | 0.504 | 0.984 | 0.996 | 0.118 |
| S×R | 8 | 0.419 | 0.009 | 0.163 | 0.089 | 0.457 |
| D×S×R | 24 | 0.998 | 0.991 | 0.994 | 0.992 | 0.100 |

D, drought intensity; S, soil substrate; R, Root order; SRL, specific root length;

SRA, specific root surface area; AD, average diameter; RTD, root tissue density

**Table S2** Carbon, nitrogen and phosphorus contents among root order of *F. mandshurica* seedlings and results of three-way (drought intensity × soil substrate × root order) ANOVA of root chemical traits.

| Root order | Soil substrate | Drought intensity | C (mg·g^-1^) | N (mg·g^-1^) | P (mg·kg^-1^) | Root order | Soil substrate | Drought intensity | C (mg·g^-1^) | N (mg·g^-1^) | P (mg·kg^-1^) |
| --- | --- | --- | --- | --- | --- | --- | --- | --- | --- | --- | --- |
| 1st | Humus | CK | 41.05±0.03a | 2.06±0.02a | 249.96±0.86a | 4th | Humus | CK | 40.99±0.03a | 2.02±0.01a | 250.85±0.80a |
|  |  | T1 | 41.05±0.05a | 2.06±0.03a | 249.55±1.37a |  |  | T1 | 40.95±0.02a | 2.05±0.01a | 252.35±0.64a |
|  |  | T2 | 41.09±0.05a | 2.08±0.03a | 248.34±1.62a |  |  | T2 | 40.92±0.02a | 1.98±0.01a | 253.26±0.66a |
|  |  | T3 | 41.01±0.04a | 2.03±0.02a | 251.04±1.35a |  |  | T3 | 40.99±0.03a | 2.01±0.01a | 250.57±1.06a |
|  | Loam | CK | 41.02±0.06a | 2.04±0.03a | 250.74±2.06a |  | Loam | CK | 40.91±0.02b | 1.98±0.01b | 253.41±0.58b |
|  |  | T1 | 41.05±0.08a | 2.07±0.04a | 249.66±2.18a |  |  | T1 | 40.88±0.04b | 1.96±0.02b | 254.37±1.38b |
|  |  | T2 | 41.01±0.04a | 2.04±0.02a | 250.91±1.28a |  |  | T2 | 41.47±0.02a | 2.06±0.01a | 281.54±0.69a |
|  |  | T3 | 40.59±0.07b | 2.02±0.04a | 217.83±2.39b |  |  | T3 | 40.95±0.02b | 1.99±0.01b | 252.12±0.79b |
|  | Sandy-Loam | CK | 41.01±0.13b | 2.02±0.06b | 251.61±4.72a |  | Sandy-Loam | CK | 40.95±0.06ab | 1.99±0.06a | 252.13±11.55a |
|  |  | T1 | 41.24±0.09ab | 2.16±0.05ab | 243.38±2.82ab |  |  | T1 | 41.01±0.04ab | 2.03±0.02a | 249.88±1.41a |
|  |  | T2 | 41.15±0.07ab | 2.11±0.03ab | 246.54±2.38ab |  |  | T2 | 40.89±0.01b | 1.97±0.05a | 254.67±0.23a |
|  |  | T3 | 41.44±0.10a | 2.26±0.05a | 237.73±2.95b |  |  | T3 | 41.04±0.05a | 2.04±0.03a | 249.42±1.69a |
| 2nd | Humus | CK | 40.97±0.03a | 2.01±0.01a | 252.25±0.91a | 5th | Humus | CK | 40.95±0.03a | 2.01±0.01a | 252.15±0.87a |
|  |  | T1 | 40.95±0.02a | 2.01±0.01a | 253.15±0.70a |  |  | T1 | 40.91±0.02a | 1.97±0.01a | 253.23±0.81a |
|  |  | T2 | 40.98±0.03a | 2.02±0.02a | 252.25±0.99a |  |  | T2 | 40.89±0.02a | 1.96±0.01a | 254.04±0.73a |
|  |  | T3 | 40.98±0.01a | 2.01±0.35a | 251.31±0.16a |  |  | T3 | 40.89±0.03a | 1.96±0.01a | 254.64±1.13a |
|  | Loam | CK | 40.97±0.01a | 2.01±0.01a | 251.96±0.14a |  | Loam | CK | 40.87±0.01a | 1.96±0.01a | 255.16±0.58a |
|  |  | T1 | 40.98±0.02a | 2.01±0.01a | 251.41±0.76a |  |  | T1 | 40.96±0.11a | 2.01±0.07a | 252.48±3.25a |
|  |  | T2 | 40.96±0.05a | 2.01±0.03a | 252.43±1.95a |  |  | T2 | 40.92±0.03a | 1.98±0.01a | 253.23±1.05a |
|  |  | T3 | 41.09±0.07a | 2.07±0.04a | 247.83±2.39a |  |  | T3 | 40.88±0.07a | 1.96±0.03a | 254.97±3.04a |
|  | Sandy-Loam | CK | 41.18±0.02a | 2.11±0.01a | 244.7±0.59a |  | Sandy-Loam | CK | 40.95±0.03ab | 1.99±0.01a | 251.87±0.95b |
|  |  | T1 | 41.11±0.07a | 2.09±0.04a | 247.57±2.36a |  |  | T1 | 40.97±0.02a | 2.01±0.01a | 250.81±0.88b |
|  |  | T2 | 41.07±0.02a | 2.06±0.01a | 248.44±0.68a |  |  | T2 | 40.89±0.02b | 1.97±0.01a | 254.69±0.78a |
|  |  | T3 | 41.13±0.05a | 2.10±0.03a | 247.01±1.77a |  |  | T3 | 40.89±0.02b | 1.97±0.01a | 254.79±0.77a |
| 3rd | Humus | CK | 41.27±0.03a | 2.07±0.01a | 282.25±0.91a |  |  |  |  |  |  |
|  |  | T1 | 40.88±0.01c | 1.96±0.25c | 254.82±0.34b | Three-ways ANOVA | | |  |  |  |
|  |  | T2 | 41.02±0.04b | 2.03±0.02ab | 249.85±1.38c | Drought intensity (D) | | | 0.338 | 0.831 | **<0.001** |
|  |  | T3 | 40.94±0.03bc | 1.99±0.02bc | 252.99±1.02bc | Soil substrate (S) | | | **<0.001** | **0.001** | **0.022** |
|  | Loam | CK | 40.90±0.01a | 1.97±0.01ab | 254.08±0.31a | Root order (R) | | | **<0.001** | **<0.001** | **<0.001** |
|  |  | T1 | 40.93±0.04a | 1.99±0.02a | 252.73±1.38a | D×S | | | **<0.001** | 0.361 | **<0.001** |
|  |  | T2 | 40.97±0.02a | 2.01±0.01a | 251.54±0.69a | D×R | | | **<0.001** | **0.044** | **<0.001** |
|  |  | T3 | 40.45±0.02b | 1.94±0.01b | 222.12±0.79b | S×R | | | **<0.001** | **0.004** | **<0.001** |
|  | Sandy-Loam | CK | 41.03±0.26a | 2.17±0.18a | 250.84±17.32a | D×S×R | | | **<0.001** | **0.045** | **<0.001** |
|  |  | T1 | 40.93±0.02a | 1.99±0.01a | 252.74±0.63a |  |  |  |  |  |  |
|  |  | T2 | 40.95±0.02a | 2.01±0.01a | 253.28±0.26a |  |  |  |  |  |  |
|  |  | T3 | 41.01±0.01a | 2.02±0.25a | 250.69±0.26a |  |  |  |  |  |  |

D, drought intensity; S, soil substrate; R, Root order;

**Table S3** Results of three-way (drought intensity × soil substrate × root order) ANOVA of roots NSCs content. Values in bold type indicate significant effects

| Source of variation | df | *P* values | | | |
| --- | --- | --- | --- | --- | --- |
|  |  | SS | ST | NSC | SS/ST |
| Drought intensity (D) | 3 | **<0.001** | **<0.001** | **<0.001** | 0.236 |
| Soil substrate (S) | 2 | **<0.001** | **<0.001** | **<0.001** | 0.075 |
| Root order (R) | 4 | **<0.001** | **<0.001** | **<0.001** | 0.440 |
| D×S | 6 | 0.743 | **<0.001** | **<0.001** | 0.310 |
| D×R | 12 | **0.011** | 0.119 | 0.115 | 0.525 |
| S×R | 8 | 0.442 | **<0.001** | **<0.001** | 0.481 |
| D×S×R | 24 | 0.269 | **<0.001** | **<0.001** | 0.568 |

D, drought intensity; S, soil substrate; R, Root order;

SS, soluble sugar; ST, starch; SS/ST, soluble sugar-starch ratio

**Table S4** Mantel analysis on the relationship between the root traits and NSC variables.

| Root order | NSC variables | *R^2^* | *P* |
| --- | --- | --- | --- |
| 1st-order | SS | 0.32 | 0.002 |
|  | ST | 0.321 | <0.001 |
|  | NSC | 0.313 | <0.001 |
|  | SS/ST | 0.238 | 0.018 |
| 2nd-order | SS | 0.278 | 0.009 |
|  | ST | 0.262 | 0.015 |
|  | NSC | 0.252 | 0.016 |
|  | SS/ST | 0.162 | 0.079 |
| 3rd-order | SS | 0.297 | 0.004 |
|  | ST | 0.136 | 0.108 |
|  | NSC | 0.118 | 0.142 |
|  | SS/ST | 0.002 | 0.892 |
| 4th-order | SS | 0.304 | 0.002 |
|  | ST | 0.136 | 0.086 |
|  | NSC | 0.115 | 0.136 |
|  | SS/ST | 0.228 | 0.01 |
| 5th-order | SS | 0.283 | 0.006 |
|  | ST | 0.068 | 0.314 |
|  | NSC | 0.042 | 0.463 |
|  | SS/ST | 0.103 | 0.16 |

SS, soluble sugar; ST, starch; SS/ST, soluble sugar-starch ratio
